# Synthesis of the novel transporter YdhC, is regulated by the YdhB transcription factor controlling adenosine and adenine uptake

**DOI:** 10.1101/2020.05.03.074617

**Authors:** Irina A. Rodionova, Ye Gao, Anand Sastry, Reo Yoo, Dmitry A. Rodionov, Milton H. Saier, Bernhard Ø. Palsson

## Abstract

The YdhB transcriptional factor, re-named here AdnB, homologous to the allantoin regulator, AllS, was shown to regulate *ydhC* gene expression in *Escherichia coli*, which is divergently transcribed from *adnB,* and this gene arrangement is conserved in many Protreobacteria. The predicted consensus DNA binding sequence for YdhB is also conserved in Entrobacterial genomes. RNA-seq data confirmed the activation predicted due to the binding of AdnB as shown by Chip-Exo results. Fluorescent polarization experiments revealed binding of YdhB to the predicted binding site upstream of *ydhC* in the presence of 0.35 mM adenine, but not in its absence. The *E. coli* MG1655, strain lacking the *ydhB* gene, showed a lower level of *ydhC* mRNA in cells grown in M9-glucose supplemented with 2 mM adenosine. Adenosine and adenine are products of purine metabolism and provide sources of ammonium for many organisms. They are utilized under nitrogen starvation conditions as single nitrogen sources. Deletion of either the *ydhC* or the *ydhB* gene leads to a substantially decreased growth rate for *E. coli* in minimal M9 medium with glycerol as the carbon source and adenosine or adenine as the single nitrogen source. The *ydhC* mutant showed increased resistance to Paromomycine, Sulfathiazole and Sulfamethohazole using Biolog plates. We provide evidence that YdhB, (a novel LysR family regulator) activates expression of the *ydhC* gene, encoding a novel adenosine/adenine transporter in *E. coli*. The YdhB binding consensus for different groups of Enterobacteria was predicted.

## Introduction

The systems biology approach for the discovery of novel gene functions, based on mRNA level analysis in *Escherichia coli* (ICA analysis) and bioinformatic approaches for the prediction of biochemical pathway regulation provide useful tools for the prediction of gene function (1,2). The inverse correlation of the transcriptional regulation of purine biosynthesis with the genes *ydhC,* encoding a putative transporter of unknown function, and *add,* encoding adenosine deaminase, suggest a relationship between YdhC, purine uptake, and purine catabolism. Growth in minimal M9 medium supplemented with adenine downregulates the PurR regulon, including the cytosine and xanthine transporter genes, *codB* and *xanP*, and cytosine deaminase, *codA*, but the *ydhC* gene, encoding a putative transporter, and the *add* gene, encoding adenosine deaminase, is upregulated (Fig. 1). As shown here, this suggestion is supported by RNA-seq analysis and metabolic pathway reconstruction. The conserved regulator gene, *ydhB,* is divergently oriented with respect to *ydhC*, logically suggesting that the former might function in the transcriptional regulation of *ydhC*.

**Figure 1.**
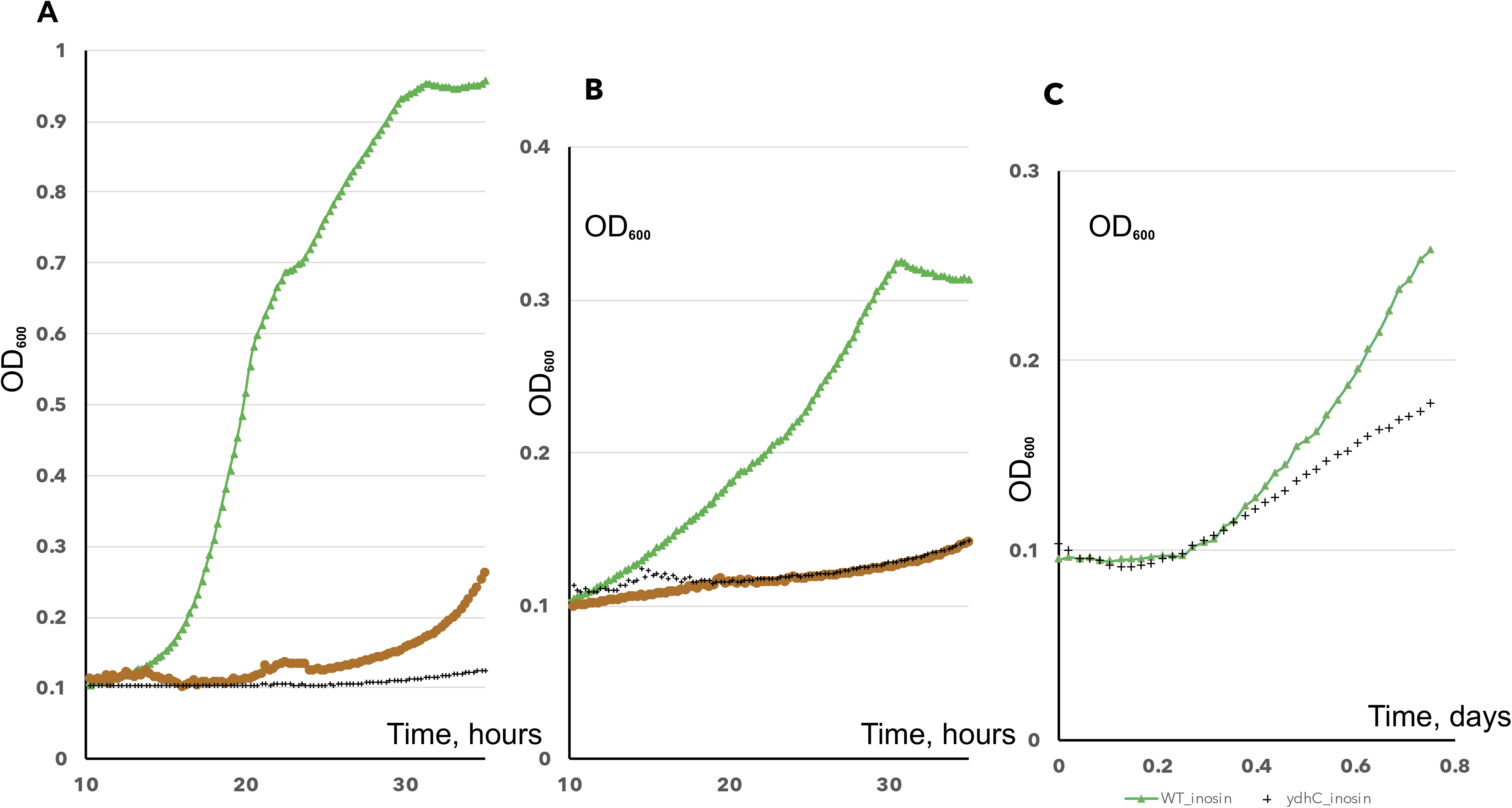
The PurR regulation of purine biosynthesis and purine/pyrimidine uptake and ydhC – add regulation **A.** The PurR i-modulone and ydhC, add genes anti-regulation. The purine biosynthesis/utilization pathways. PurR regulated genes are shown in green and blue boxes. Abbreviations: XanP, CodB – xanthine and cytosine transporters, CodA -cytosine deaminase, Add - adenosine deaminase, AdeD - adenine deaminase, YdhC – novel adenosine/inosine transporer, PpnP – adenosine phosphorylase. IMP - inosine monophosphate. The PurFDTLMEKCBH enzymes (IMP biosynthesis) encoding genes and xanP are regulated by the PurR and anticorrelated with ydhC-add transcription. **B.** The binding of YdhB to the ydhC promoter, shown by ChIP-Exo and YdhB Logo for the binding site. **C.**

Adenosine (Adn) transporters are important as mediators of the uptake of Adn when present in the medium as a substitute for a more traditional purine, carbon or nitrogen source. NupC and NupG are previously characterized transporters in *E. coli* for nucleosides when used as a carbon source. They recognize the nucleoside ribose moiety (3) and are regulated by the carbon catabolite repressor protein, CRP. NupC and NupG were shown recently to be important ADP-glucose uptake porters and these transporters are essential for the incorporation of extracellular ADP-glucose into glycogen during its biosynthesis (4). The coordinate regulation of different purine transporters is important for the utilization of nucleosides as carbon and nitrogen sources under various starvation conditions (5).

Adenosine is utilized via adenosine deaminase (Add) which converts it to inosine. PpnP is a broad specificity pyrimidine/purine nucleoside phosphorylase that produce hypoxanthine (Hpx) (adenine) and D-ribose-1-phoshate from phosphate and inosine (adenosine) respectively. The PurR regulon is essential for purine biosynthesis, and PurR is activated by the presence of hypoxanthine (Hpx), signalling that the purine concentration is sufficient for growth (5,6). The inverse correlation for the modulation of mRNA levels for PurR-regulated purino biosynthetic genes and those of the *ydhC* and *add* genes (7) has been demonstrated during growth in minimal M9 medium with adenine as a supplement (1). For PurR regulon related genes, the levels of mRNAs encoding proteins of purine/pyrimidine biosynthesis and guanosine uptake decrease in the presence of adenine, while *ydhC* and *add* gene expression increases (1,6). Thus, purine biosynthesis pathway genes, *purBCDEFHLMN*, are repressed by PurR in response to the availability of cellular hypoxanthine as a product of adenosine degradation or purine biosynthesis. Extracellular adenosine gives rise to hypoxanthine due to uptake followed by the action of the *add*-encoded enzyme, adenosine deaminase, and further, by the PpnP (YaiG) catalyzed reaction (Fig. 1). Adenine uptake as a nitrogen source or purine precursor leads to an increase of Hpx as a product of the AdeD (adenine deaminase) catalyzed reaction.

We hypothesize that YdhC is an adenosine transporter and that YdhB is the transcriptional activator for the *ydhC* gene. The poor fitness phenotype for a *ydhC* homolog mutant in *Pseudomonas simiae WCS417* (66% identity with the *E.coli* protein) has been shown during growth in minimal medium with adenine as the nitrogen source. It was mildly important with adenosine as the nitrogen source (fit.genomics.lbl.gov). This hypothesis is supported by the shown strong fitness phenotype of *Klebsiella michiganensis M5al* for the *ydhC* mutant homolog (75% identity with the *E. coli* homolog). This mutant was found to have a strong negative fitness during growth with any one of several purines as carbon sources: inosine, 2-deoxy-inosine, 2-deoxy-adenosine, 2-deoxy-adenosine 5-phosphate as well as a mild negative effect with adenosine as the carbon source.

## Results

The work described here expands our understanding of how adenosine uptake and degradation versus biosynthesis are reciprocally regulated. Under the conditions described here, extracellular adenosine or adenine is the only nitrogen source supporting the growth of *E. coli*, but it does not support growth of a *ydhC* deletion mutant strain. The hypoxanthine produced from adenosine is a signal for the PurR-mediated repression of purine biosynthesis. This proposed regulation of the PurR modulon is shown in Fig. 1.

### Prediction of YdhC function based on the regulatory modulation by PurR

The independent component analysis (ICA) revealed regulation of the PurR modulon, consisting of the *purABCDE,* genes and those encoding the transporters for xanthosine (XanP) and cytosine (CodB) as well as the inverse regulation of the *ydhC* and *add* genes in response to the presence of adenine in the growth medium (1). We hypothesized that YdhC is an adenosine/adenine transporter because the presence of a purine source in the medium should inhibit biosynthesis, mediated by the PurR-hypoxanthine complex while upregulating the adenosine/adenine transporter. Although NupC and NupG are known nucleoside transporters, regulation by CRP suggest that they are essential for nucleoside uptake to be used as carbon sources. NupG requiress hydroxyl groups in the ribose moiety for substrate binding.

YdhB is homologous to the AllS transcriptional regulator (HTH-type) of the LysR family. AIIS activate as the allantoin utilization cluster in *E. coli*.(Ref) The *ydhB* and *ydhC* genes are divergently transcribed from each other, and this arrangement is conserved in other Proteobacteria (Fig. 2A) (see Introduction). Alignment of the upstream regions for the *ydhC* genes revealed a palindromic consensus for binding of the predicted YdhB regulator (Fig. 2B). A phylogenetic footprinting approach was used to define the level of conservation of a AdnB⎕protected region from *E. coli* in other gamma-proteobacteria possessing AdnB orthologs. In addition to various species from the Enterobacteriaceae family, *adnB (ydhB)* orthologs were identified in *Pseudomonas* spp., and in all cases *ydhC* orthologs are located immediately upstream of *ydhB* with a shared promoter region. For each subset of closely-related enterobacteria and *Pseudomonas* we generated multiple sequence alignments of orthologous *ydhB* upstream regions. The AdnB⎕protected region from *E. coli* corresponds to a highly conserved fragment of these regions (Fig. S2). By combining these orthologous AdnB⎕protected regions we defined the conserved 23-bp AdnB⎕binding motifs for Enterobacteria and *Pseudomonas* spp. (see sequence logos in Fig. S2). A common motif of these taxonomy-specific AdnB⎕binding sites is an imperfect palindrom with consensus TsttwTCAAwAwwwTTGaaGGCA, where ‘s’ is either G or C and ‘w’ is either A or T.

**Figure 2.**
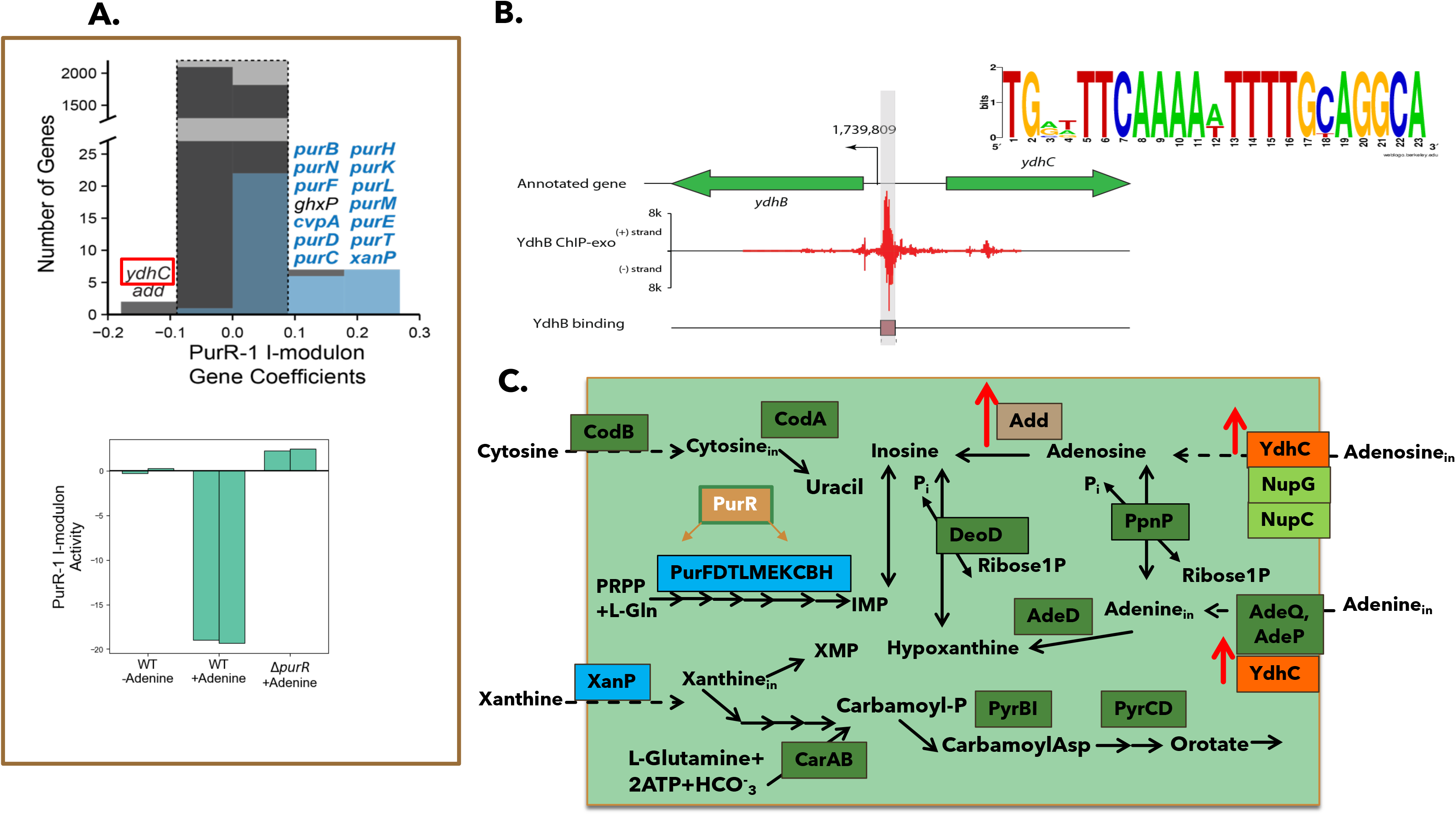
The growth of the ydhB (circle), ydhC (plus) mutants compared to the WT BW25113 (triangle) strain in M9 medium with 2.5 mM adenosine or inosine as the sole nitrogen source and glycerol or glucose as the carbon source, **A.** M9 with glucose as carbon source and adenosine as sole nitrogen source **B.** M9 with 0.4% glycerol as carbon source and adenosine as nitrogen/carbon source **C.** M9 with 0.4% glycerol as carbon source and 20 mM inosine/glutamate as nitrogen sources.

### Growth in M9 medium with adenosine as a supplement

RNAseq analysis of the *ydhB* mutant of *E. coli* MG1655 (WT) in M9 medium supplemented with 2.5 mM adenosine showed downregulation of the *ydhC* gene, suggesting that YdhB is an activator for *ydhC* expression. Supplementation of M9 medium with adenosine increased *ydhC* mRNA only in the presence, but not in the absence, of YdhB. ICA analysis compared the modulons for growth in M9 medium with adenosine supplementation as the sole nitrogen source. The modulon regulated by FliA was downregulated in the WT strain, but not in the *ydhB* mutant strain. FliA regulates the motility in *E. coli,* and the FlhDC modulon was dowregulated in both strains (Ref).

### Growth in M9 medium with adenosine as nitrogen and glycerol as carbon sources

When adenosine was added to the medium under nitrogen limiting conditions, the growth of *E. coli* BW25113 was supported, but no growth was observed for the *ydhC* mutant (Fig.3), either with glycerol or with glucose as the carbon source. The effects on growth with adenosine for the *ydhB* mutant is more substantial with glycerol as the carbon source; the growth effects are shown in Figs. 3A and B. We suggest that high concentrations of adenosine can support metabolism via the pentose phosphate pathway, producing ribose phosphate.

### Growth in M9 medium with inosine and glutamate as sole nitrogen sources and glucose as the carbon source

The supplementation of M9 minimal medium with inosine in addition to glutamate was tested for the WT and *ydhC* mutant strains for growth with glycerol as the carbon source. We also compared growth in M9-glucose medium with inosine/Glu as nitrogen sources. The *ydhC* mutant showed a decrease in the growth rate compare to the WT strain under nitrogen starvation conditions using inosine as an additional nitrogen source (Fig. 4).

### YdhC specificity screening for carbon sources using Biolog plate 1

The growth rates for the *ydhC* mutant and wild type *E. coli* BW25113 (WT) using different carbon sources, including adenosine or inosine on Biolog plate 1 was measured using M9 minimal medium without another carbon source. The strains were grown overnight in LB and washed twice with M9 medium without addition of a carbon source. The cultures were diluted, and 0.1 ml of each was added to the 96-well plates. Growth was detected with Omnilog, but no difference for adenosine or other carbon sources were detected for the mutant and WT strain under microaerobic conditions.

### Fluorescent polarization (FP) assay for AdnB binding to the DNA

The purified YdhB_His protein was incubated with a fluorescently labelled DNA fragment, predicted by phylogenetic footprinting. YdhB protein (0 - 60 nM) was incubated in the assay mixture described in Materials and Methods with a 10 mM MgSO_4_ supplement for 1 hour at room temperature. The FP signal at different concentration of AdnB is shown in Fig. 3. The AdnB binding in the presence, but not the absence of 0.35 mM of adenine was detected using the fluorescence polarization assay. The AdnB concentration K_d_ = 27nM for binding to DNA was calculated using Prism 7.

### YdhC specificity screening for antibiotics using Biolog plates 11C and 12

The differences in growth rate for the *ydhC* mutant (from the Keio collection) compared to wild type *E. coli* BW25113 (WT) were measured using Biolog plates 11C and 12 for aminoglycosides. Biolog plates 11C and 12, contain 4 different concentrations of 24 different antibiotics (Fig.3). The *ydhC* and WT strains showed the same resistance for all of them. However, the mutant strain showed increased resistance to paromomycine as well as sulfathiazole and sulfamethoxazole compared to the WT *E. coli* strain. Paromomycine is an aminoglycoside group antibiotic, inhibiting protein synthesis. The increased resistance observed for the YdhC transporter mutant strain suggests specificity for uptake of paromomycine, sulfamethoxazole and sulfothiazole. The last two antibiotics are structural analogs of para-aminobenzoic acid (PABA). They compete with PABA for binding to dihydropteroate synthase and inhibit conversion of PABA and dihydropteroate diphosphate to dihydrofolate. It is interesting that a distant *ydhC* paralogous gene in *E. coli* encodes Bcr – a bicyclomycin/sulfonamide resistance protein (8). Bcr has been shown to be involved in the export of L-cysteine (9). In contrast, YdhC seems to be an adenine/adenosine uptake transporter with broad specificity, and it may also be involved in sulfamethoxazole and sulfothiazole, uptake both sulfonamide group antibiotics, and paromomycin uptake.

## Materials and methods

### Constructs of the *adnB (ydhB)* and *adnC (ydhC)* mutants, and overproduction of AdnB (YdhB)

The *ydhC* and *ydhB* mutants from the Keio collection (10) were grown overnight in LB medium. Then the cells were refreshed for 3 hours, washed with M9 salts medium and inoculated in the M9 medium lacking the usual a nitrogen source, but substituting it with 2.5 mM adenosine or 2.5 mM adenine. The *adnB* gene sequence was confirmed by re-sequencing it using ASKA collection primers.

The *adnB* overexpressing strain was inoculated from the ASKA collection (11) onto LB agar plates containing chloramphenicol. Overnight cultures were then inoculated from single colonies. 50 ml of each of the new cultures was started, and after the OD_600_ reached 0.8, 0.8 mM IPTG was added. The cultures were incubated at 24°C overnight with continuous shaking, and cells were collected by centrifugation.

The AdnB recombinant protein, containing an N-terminal 6His tag, was purified by Ni-chelation chromatography from the soluble fraction as described (12,13). The insoluble fraction was solubilized in 8M urea and purified on a Ni-NTA minicolumn with At-buffer (50mM Tris-HCl buffer, pH 8, 0.5 mM NaCl, 5 mM imidazole, and 0.3% Brij) with 7M urea. The purification procedure has been described in detail (12).

### The fluorescence assay

The AdnB binding assay mixture (0.1 ml) contained Tris buffer, pH 7.5, 0.1 M NaCl, 0.5 mM EDTA, 10 mM MgSO_4_, 2 mM DTT, 5 μg/ml sperm DNA and 1μM of the fluorescently labelled predicted AdnB binding DNA fragment as well as 0-0.6 mM adenine or 3 mM AMP. Then the AdnB protein (0-1 μM) was added to the assay mixture, and it was incubated for 1 hour at 30°C.

### Prediction of the regulatory binding site

A bioinformatic approach to phylogenetic footprinting described in (14) was applied for the prediction of the DNA binding site in the intergenic region of *adnB-ydhC*. The intergenic region was aligned for different representative Proteobacteria from the genomes listed in Table 1. Conserved nucleotides were predicted using MEME program.

## DISCUSSION

The presence of adenosine can support *E. coli* growth as the sole nitrogen source, and it can be involved in acid resistance (15). Additionally, inosine or guanosine in the *E. coli* growth minimal medium can increase the growth rate as does arginine, but they cannot support growth as the sole nitrogen source. The adenosine degradation pathway involves Add (adenosine deaminase) to produce inosine. Deletion of *add* attenuates growth in the presence of adenosine under acidic conditions (15). We found that YdhC is essential for adenosine and adenine uptake under nitrogen starvation conditions, and it could be involved in acid resistance in the presence of adenosine. However, under aerobic growth at pH 5.5 with adenosine or inosine as a supplement, it has not been shown to have a positive effect on acid resistance in the BW25113 strain (Fig. S1).

The transporter, YdhC, and the *ydhC* gene regulator, YdhB, are conserved as is the *purR* gene (Fig. S2) in many proteobacterial genomes. The HTH-type regulator, YdhB, homologous to AllS (Fig. 5), was shown to be a transcriptional activator for *ydhC* gene expression. YdhB is also essential for growth during nitrogen starvation with adenosine as the sole nitrogen source and glycerol as the carbon source (Fig. 2). The pathway for adenosine degradation probably supports carbon flux to ribose-phosphate through the pentose-phosphate pathway. The pathway for purine biosynthesis, uptake and degradation is shown at Fig. 1. It is interesting that previously, AdhB and YdhC had been shown to be essential for *Yersinia pseudotuberculosis* growth, suggesting that adenosine autotrophy may be a property of many Proteobacteria as also suggested from the genome analyses.

The potential specificity of *ydhC* for the uptake of certain aminoglycosides such as paromomycine-related antibiotics was demonstrated. The *ydhC* strain was shown to have substantially increased resistance to paromomycine, although it showed elevated resistance to sulfathiazole and sulfamethoxazole compared to BW25113 WT E. *coli*

We found that a palindromic sequence between *ydhB* and *ydhC* serves as the YdhB binding site, conserved in many Proteobacteria. The binding was shown for *E. coli* using ChIP-Exo and a fluorescent polarization assay. The gene, *ydhB*, in many Proteobacterial genomes, including all Enterobacteria, together with the *ydhC*, and the *purR* genes is conserved in the same gene contexts. All of these observations substantiate the main conclusion of this paper.

## Acknowledgements

This work was supported by National Institutes of Health (NIH) grant U01AI124316 and Novo Nordisk Foundation Grant Number NNF10CC1016517 as well as NIH grant GM077402.

