## Supplemental Fig 2 for "Synthesis of the novel transporter YdhC, is regulated by the YdhB transcription factor controlling adenosine and adenine uptake"

Enterobacteria -1

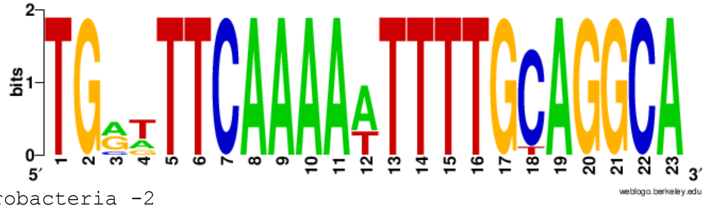


Enterobacteria -2

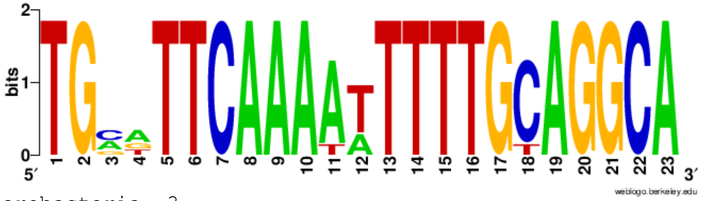


Enterobacteria -3

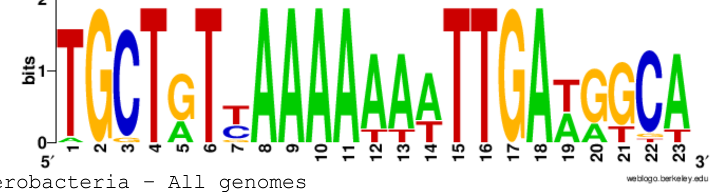


Enterobacteria - All genomes

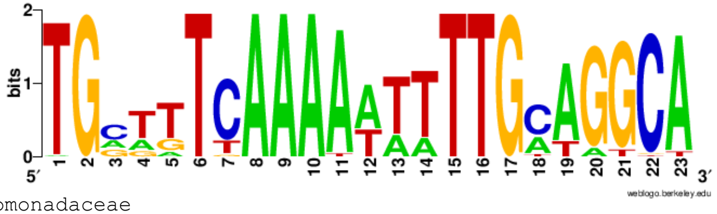


Pseudomonadaceae

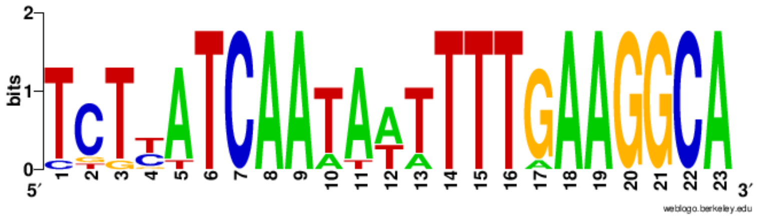
