## Supplemental FigS1 for "Synthesis of the novel transporter YdhC, is regulated by the YdhB transcription factor controlling adenosine and adenine uptake"

### Slide 1
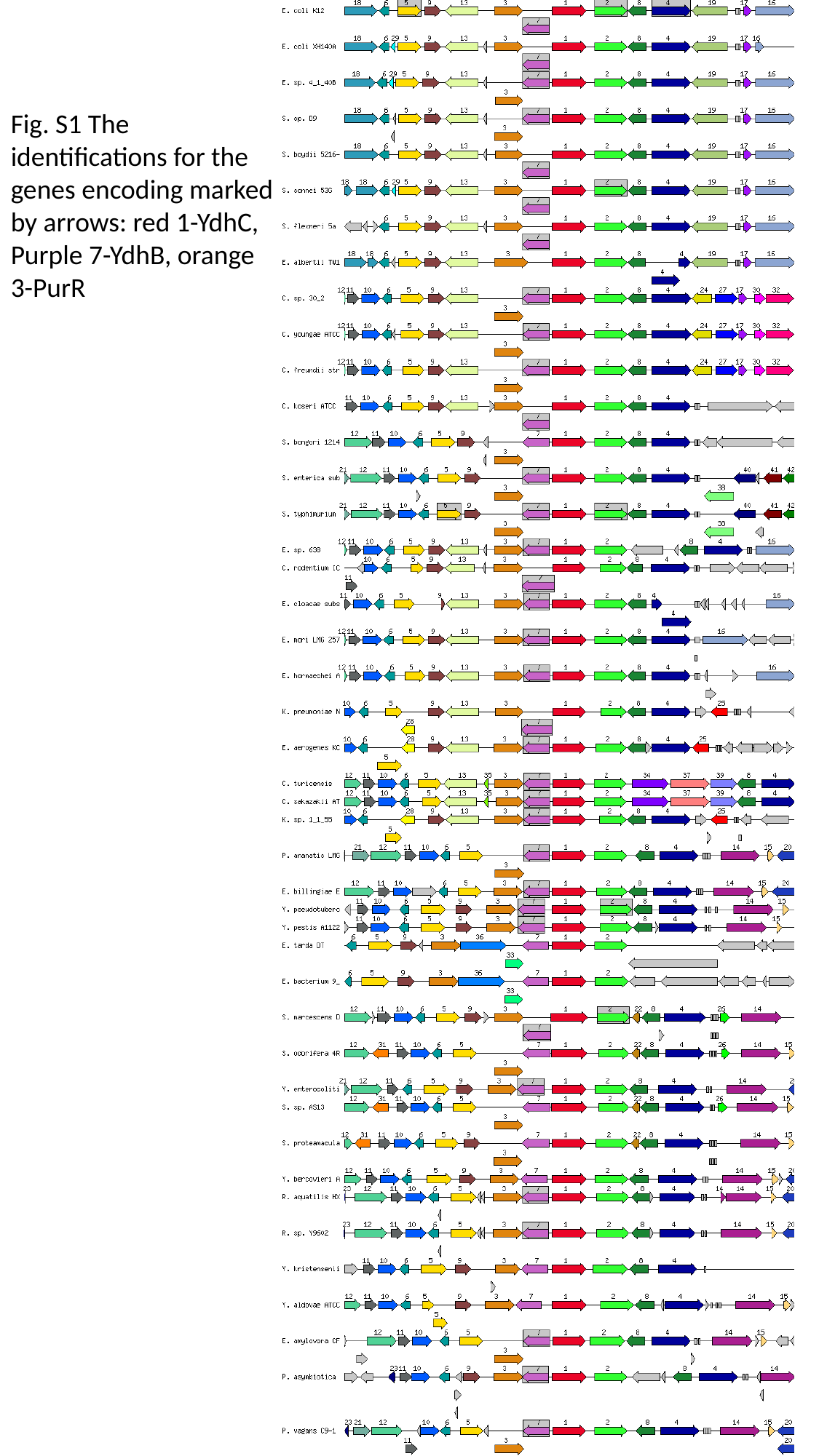

Fig. S1 The identifications for the genes encoding marked by arrows: red 1-YdhC,
Purple 7-YdhB, orange 3-PurR
